# Root and shoot competition lead to contrasting competitive outcomes under water stress: A Meta-analysis

**DOI:** 10.1101/712208

**Authors:** Alicia J. Foxx, Florian Fort

**Author notes:** to whom correspondences should be addressed. (AF). Author conceptualized the study, performed analyses and visualizations, wrote manuscript (Alicia J. Foxx). Author reviewed and edited this manuscript (Florian Fort).

## Abstract

**Background:** Competition is a critical process that shapes plant communities and interacts with environmental constraints. Though important to natural communities and agricultural systems, there are surprising knowledge gaps related to mechanisms that belie those processes: the contribution of different plant parts on competitive outcomes and the effect of environmental constraints on these contributions.

**Objective:** Studies that partition competition into root-only and shoot-only interactions assess whether plant parts impose different competitive intensities using physical partitions and serve as an important way to fill knowledge gaps. Given predicted drought escalation due to climate change, we focused meta-analytic techniques on the effects of water supply and competitive outcomes.

**Methods:** We searched Web of Science for peer-reviewed studies and found 2042 results. From which six suitable studies with 92 effect sizes on 10 species were identified to test these effects.

**Results:** Water availability and competition treatment (root-only, shoot-only, and full plant competition) significantly interact to affect plant growth responses (p < 0.0001). Root-only and full plant competition are more intense in low water availability conditions than shoot-only competition. Shoot-only competition in high-water availability was the most intense showing the opposite pattern. These results also show that the intensity of full competition is similar to root-only competition and that low-water availability intensifies root competition while weakening shoot competition.

**Conclusions:** These results emphasize the importance of root competition and these patterns of competition may shift in a changing climate, creating further urgency for further filling knowledge gaps to address issues of drought on plant interactions and communities.

## Introduction

A major question among plant ecologists is to understand plant competition mechanisms and their outcomes from different perspectives. Many contemporary ecological endeavors seek to elucidate the role of competition in community structure, processes, and species coexistence [1–6]. Evidence shows that competition impacts survival, and higher level processes such as community diversity and spatial structure [7,8] Past work dived deeply into understanding the role of pair-wise species competition on outcomes observed in communities and in field settings [9–12]. But, only a small section of the literature describes the competitive contributions of roots and shoots separately (Fig. 1) and their interaction with environmental constraints - which is critical considering the contribution of roots and shoots to ecosystem processes and responses to environmental changes[13–15].

**Figure 1.**
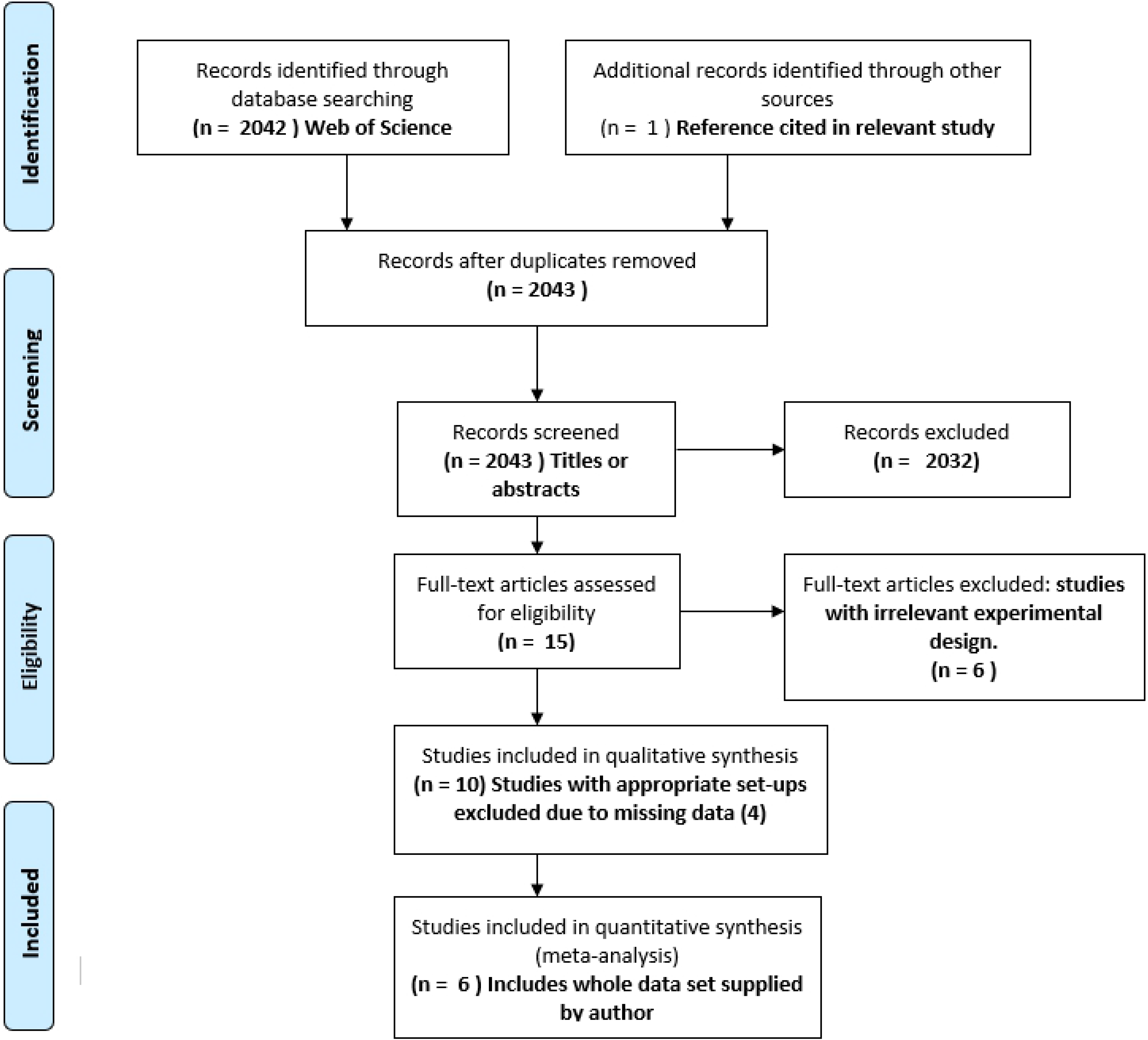
Study treatments. Competition treatments of root-only, shoot-only, full competition and, monoculture of partition studies.

Most competition studies focus on competitive outcomes on shoots. But competitive behaviors resulting from shoot competition, may not influence competitive root responses in the same plant [16], thus the influence and outcome of roots interaction needs specific consideration. Traits can predict competitive ability and performance in environments [17,18], and Kembel & Cahill [19] showed that roots face different environments than shoots leading to variable correlation of above- and belowground traits in response to the environment. A meta-analysis on studies that physically partitioned roots and shoots during competition under nutrient stress found that roots imposed more intense competition than shoots reporting a 42% biomass reduction – indicating intense competition. [20]. A critical remaining question is on the role of water in competition.

Water is a critical resource that allows plant growth, and related physiological processes such as cell growth and nutrient transport to shoots [21,22]. In case of low water availability plants can close stomata to limit water loss and CO_2_ capture [23]. They can also respond to water stress by allocating more mass to roots to acquire the limited resource [24,25]. Generally, while water stress reduces plant size, root allocation, branching, length, and uptake, increase to maintain soil water capture capacities [26–29] (Fig. 2). Conversely, water stress reduces shoot growth, leaf area, new leaf production, and photosynthetic light conversion [27,29–31] (Fig. 2). Resulting diminished light interception and metabolic activity aboveground [32], coupled with increased absorptive root area under water stress should intensify competition between roots more than between shoots (e.g. [33]), but the literature presents mixed evidence related to their outcomes.

**Figure 2.**
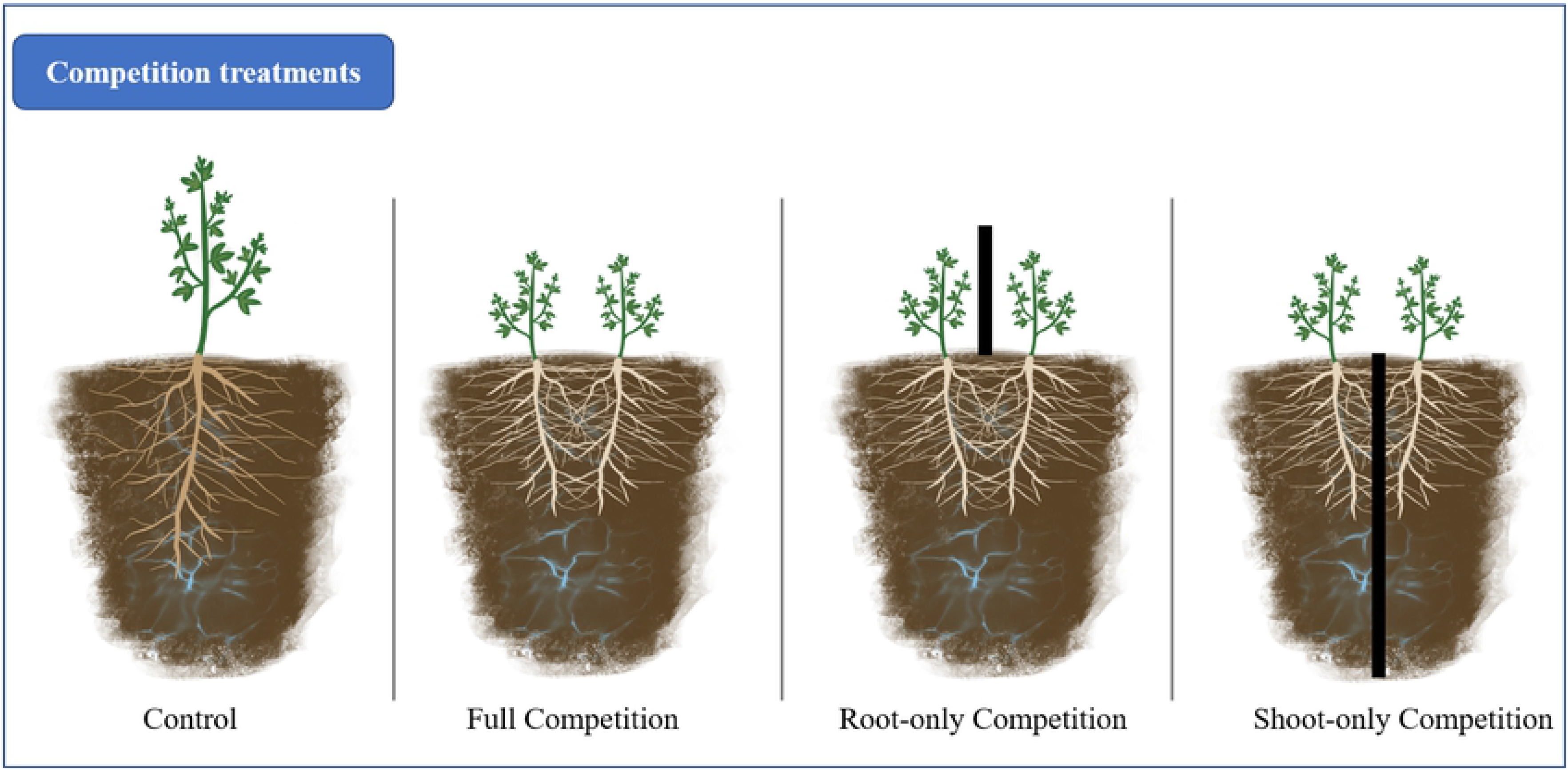
Competition and water stress impacts. Morphological and physiological above- and belowground competitive responses to water availability.

Despite established patterns of individual effects of water stress, water stress intensifies, decreases or produces no measured outcomes on root-only or shoot-only competition (e.g. [34–36]. The different physiological processes of roots and shoots to drought, may reduce resource acquisition need. These differing activity levels during drought may also have strong effects on above- compared to belowground performance that may affect the intensity of root and shoot competition in water limited environments. This is critical due to the predicted variable global precipitation patterns and increased regional aridity due to climate change [37]. Environmental constraints such as resource stress change the intensity of the competition among species [38–41]. For example, low water availability can intensify [42,43] or weaken competition [44] and, for example, water loss of a nurse shrub due to dry soil reduced mortality in a protégé shrub [45]. Despite the substantial impacts water limitation imposes on competition and survival compared to nutrient stress [46], the literature pool on water and competition is comparatively small so synthesis would advance our knowledge by elucidating patterns.

We conducted a meta-analysis to provide resolution on the intensity of root and shoot competition under water stress. We assessed whether roots and shoots impose different competitive intensities in studies that physically partitioning roots and shoots during competition experiments under different water availabilities (Fig. 1). We hypothesize that: 1) competitive intensity of root-only, shoot-only, and full competition will differ under varying water availability; 2) competitive intensity will differ between low – and high-water stress treatments; and 3) root competition will differ from shoot competition at varying water availabilities.

## Methods

### Literature Search

We sought peer reviewed literature using the Web of Science searching platform. A search was performed on 2 May 2019 of the following title and topic with Boolean terms and wildcard symbols to broaden the search: [(shoot* AND root*) OR (above AND below)] AND [(competit* OR interact*)], topic: “water stress.” Search results were refined by research areas of plant sciences, agriculture, genetics, heredity, forestry, and environmental sciences, and ecology. See Supplemental figure S1 for study selection flow diagram (PRISMA checklist, S1 Table 2). Citations within relevant articles were searched as well. Abstracts were then evaluated for relevance and kept if they met the following experimental criteria: experimental designs that contained root-only, shoot-only, and or full competition, and a control group (Fig. 1), all under a high- and low-water availability treatments. Weigelt et al. [36] lacked a shoot competition treatment but was included here. Authors were contacted for data sharing when standard deviation, sample size, and/or response variables were not reported, imputable, or extractable.

### Data Collection

Studies were included in the analyses if we acquired response variables, standard deviation, and sample sizes, either from the study, the study authors, or from figures. When data were only available in graphics, those data were extracted from figures using the free web-based application WebPlotDigitizer v4.1 [47]. We extracted data from figures from three studies [34,48,49]. Two studies implemented multiple water treatments [50,51], so data from the two extreme treatments were used (highest and lowest water availability). Nutrient treatments were used in some studies but this was not replicated in all studies so, only data from the lowest nutrient level were utilized. Fixed effects from each study included water treatment (low- and high-water availability treatments), competition treatment (control, root-only, shoot-only, or full competition) (Fig. 2), and focal species nested within study as a random effect.

### Analyses

We constructed mixed effects meta-regression models to compare the log response ratio values. Models were constructed using the “rma.mv” function in the “metafor” package (Viechtbauer 2010) in R [52]. Models were compared using logliklihood ratio test that used the “anova” function. To test whether water treatments modulated outcomes of the competition treatments, the full model assessed the interaction between water availability levels (low- and high-availability) and competition treatments (root-only, shoot-only, and full). The reduced models were compared to the full model to determine which explained more variation in plant growth. The reduced models assessed plant growth response to water availability, or plant growth responses to competition treatment, and plant growth responses to the additive effects of competition and water treatments.

The effect sizes of the response variables were calculated as log response ratios [lnRR; 53]. Log response ratios are the proportional change in treatment groups compared to the control group [53]. They are symmetric around zero and taking the log linearizes the ratio and leads to a generally normal distribution when the treatment mean is not zero [53]. The lnRR values are approximately normally distributed [53]. Negative values denote intense competition and positive values denote weak competition while a lnRR of zero denotes no effect of treatment [54]. The lnRR values were calculated in R using the “ROM” measure in the “escalc” function in the “metafor” package [55]. The “ROM” measure belies the equation:

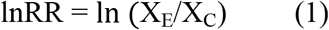

Where X_E_ is the response means of plants under root-only, shoot-only, or full competition compared to the mean of the control group X_C_. The lnRR values were calculated over study and species and compared between root-only, shoot-only, and full competition, as well as water availability levels. The calculated effect size is the most likely effect size but confidence intervals are important in interpreting meta-analyses outcomes [54]. They indicate how confident one is in the directionality of an effect size and tell the full range of effect size for the treatment [54](Valentine et al. 2010). If the lower bound confidence interval overlaps with zero, the results are not statistically significant (Valentine et al. 2010). The sampling variances of the lnRR were calculated in R using the “escalc” function, and the equation follows Hedges et al. [53]: Where n_E_ and n_C_ and SD_E_ and SD_C_ are the sample sizes and standard deviations for the experimental and control groups respectively. Standard deviation was not reported in three suitable studies [48,50,51], but standard deviations were imputed to reduce publication bias and improve variance estimates compared to when data from an incomplete study are excluded [56]. So, the standard deviation from one study [51] was calculated using F-statistics reported in the original study using equation 3 (Larry Hedges, Personal communication):

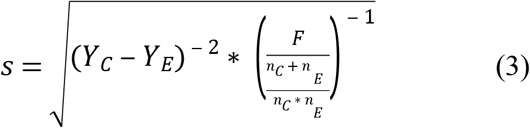

where n_C_ and n_E_ the sample sizes of the control group and treatment group respectively. Additionally, Y_C_ and Y_E_ are the mean values of the control group and treatment group respectively, and s is the standard error. Standard deviations were also imputed for two studies [48,50] using a linear regression between sample sizes and pooled standard deviation values of studies with known standard deviation values using the following equation [56]:

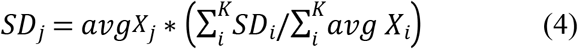

where s_j_ is the standard deviation of the study with missing information and SD_i_ is the standard deviation of samples with full information, X_i_ is the mean of the LRRs of full studies and Xj is the mean of the LRRs of the study with missing information.

We performed contrasts to test the hypotheses that root competition differed from shoot competition at differing water levels, and the hypothesis that competitive intensity differed between water availability levels. Contrasts were specified in the “linearHypothesis” function in the “car” package [57]. All analyses were performed in R [52].

**Table 1.** Characteristics of studies used for meta-analysis

| Study | Experimental Setting | Target Species | Response measure |
| --- | --- | --- | --- |
| Bartelheimer et al. 2010 | Outdoor - mesocosm | *Senecio aquaticus*; *Senecio jacobaea* | Total biomass |
| Bornkamm et al. 1975 | Setting unknown – pots | *Arrhenatherum elatius*; *Bromus erectus* | Root biomass |
| Lamb et al. 2007 | Outdoor – plots | *Artemisia frigida*; *Chenopodium leptophyllum* | Shoot biomass |
| Salinger & Bornkamm 1982 | Outdoor – pots | *Arrhenatherum elatius*; *Bromus erectus* | Shoot to root ratio |
| Weigelt et al. 2005 | Outdoor - mesocosm | *Carex arenaria*; *Corynephorus canescens*; *Hieracium pilosella* | Total biomass |
| Wilkinson & Gross 1964 | Greenhouse – pots | *Trifolium repens* | Total biomass |

## Results

### Literature search

The search results yielded 2042 studies (SI Fig. 1). Eleven studies with appropriate methods were found. One researcher provided data from her study (Weigelt et al. 2005). Four were excluded as data useful for calculating effect sizes and variance were unavailable in figures or through authors. One excluded study used trees as focal plants [58] while all others utilized herbaceous or shrub species. Furthermore, this study [58] and another [35– also with missing data] imposed water availability treatments by differences in water availability by site [58]. This introduces large heterogeneity and doubts on whether the effect sizes are drawn from the same population – an assumption of fixed effects meta-analytic models [59]. This left six studies with extractable information. These six studies on ten species contained 130 data points, and 92 lnRR outcome measures (Data for calculations; SI Table 1).

### Competitive outcomes

The model that best fit the data included an interaction between competition and water treatments (Q_df=5_ = 32.7, p < 0.001) (Table 2), whereby competition and water treatments interacted to significantly affect plant growth. Root-only, shoot-only and full competition exhibited different responses to water treatments (Fig. 3, Table 2). Root-only (−39%) and full (−13%) competition at low water availability was more intense than shoot-only (+54%) competition. Root–only (+27%) and full (+17%) competition under low-availability were more intense than shoot-only competition (−68%).

**Table 2.** Table of model outcomes. Q test statistic assess significance of between study variation [53]; T^2^ measures the between study variance; I^2^ measures variance explained by heterogeneity between studies [55]; and sigma denotes the between study variance component [53].

| Factor | Df | Q | $T^2$ | $I^2$ | P -value |
| --- | --- | --- | --- | --- | --- |
| Competition treatment | 1 | 32.3 | 0.54 | 98.9% | <0.0001 |
| Water treatment | 2 | 192.9 | 0.57 | 99.4% | <0.0001 |
| Competition + water treatment | 3 | 31.15 | 0.53 | 98.7% | <0.0001 |
| Competition * water treatment | 5 | 362.5 | 0.53 | 98.1% | <0.0001 |
| Random effects<br>(study/species) | Sigma = 0.13 |  |  |  |  |

**Figure 3.**
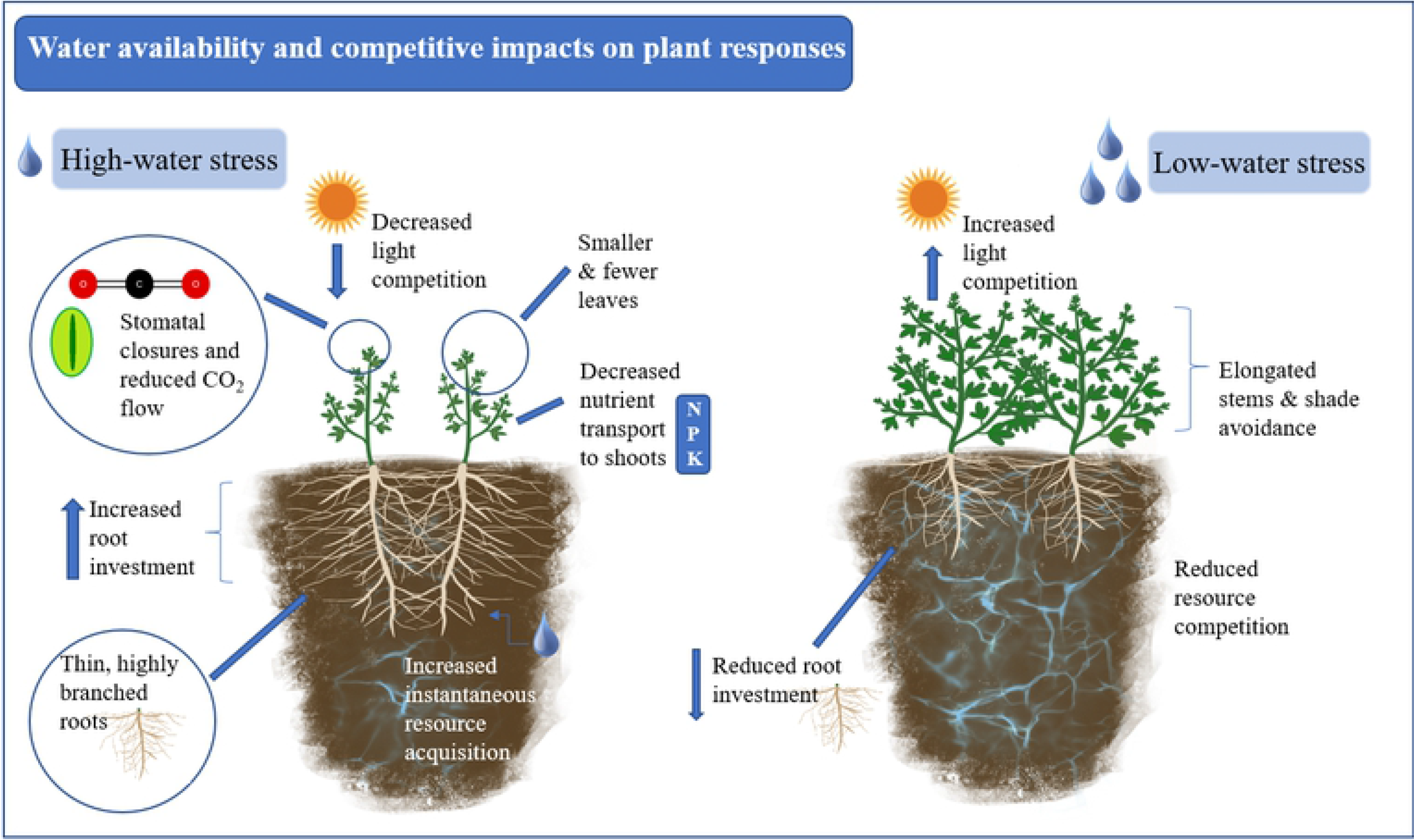
Effects of water availability and competition on plant growth. Meta-estimates (diamond points) and 95% confidence intervals. Smaller values indicate intense competition, while larger values indicate weaker competition. Percent values denote treatment effect compared to the control group: smaller (−) or larger (+) (lnRR * 100%). Number of effect sizes is in parentheses.

**Figure 4.**
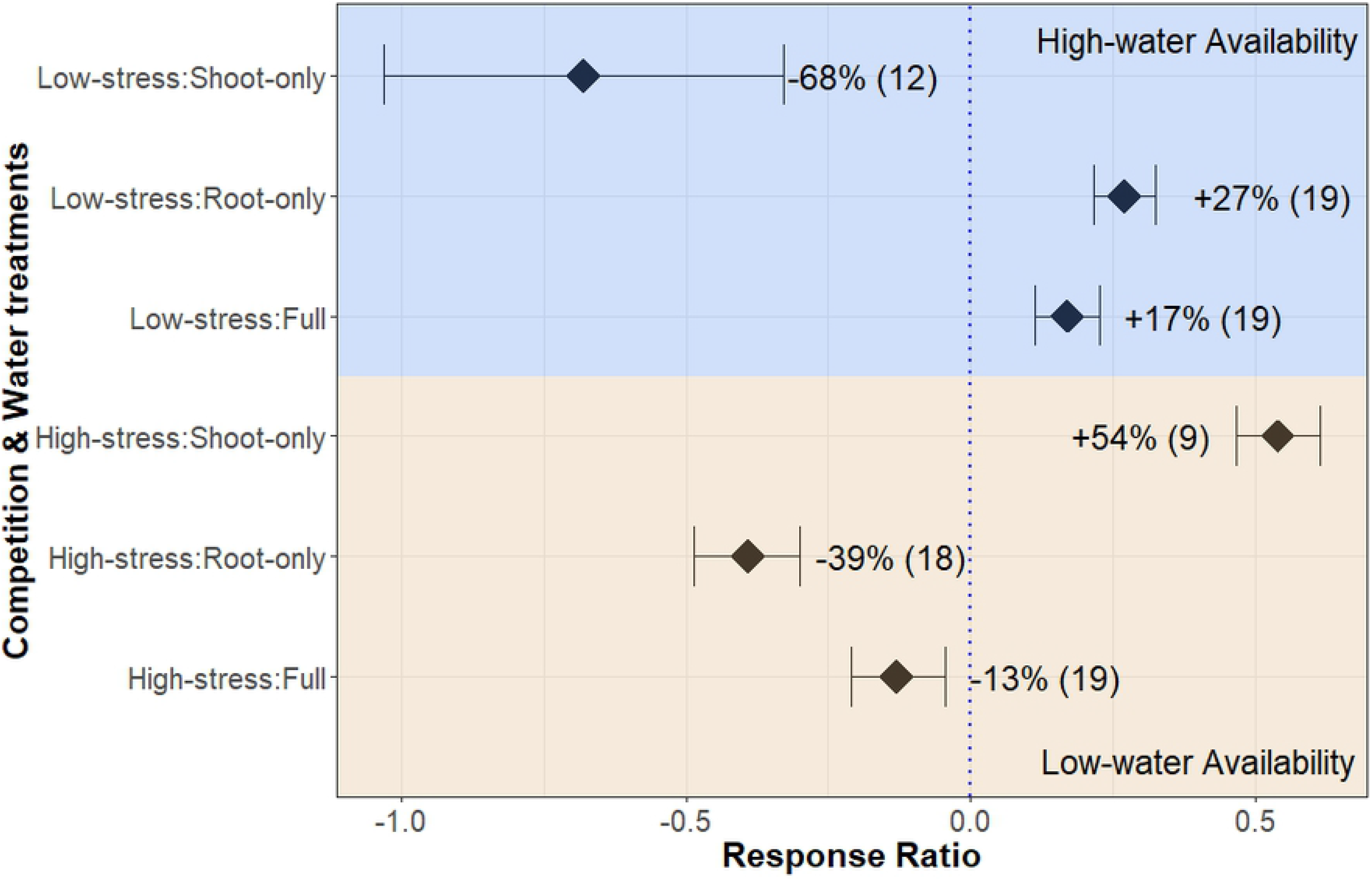

The heterogeneity between studies (Q_df=84_) is 782.1 indicating that heterogeneity between studies is high (given a Q > 100 we reject the null hypothesis that the variance component is 0 [53]) and there are likely differences between studies and unexplored sources of variation we did not capture. This is reinforced by the high I^2^ values (Table 2) denoting that a large part of the variation remains unexplained. Root-only and shoot-only competition had significantly different responses to water treatments (p <0.001) where root-only competition was more intense than shoot-only competition under low-waters availability and the opposite pattern at high water availability treatments (Fig. 3).The overall plant response was not significantly impacted by water availability (p = 0.1). Low-water availability caused slightly weaker competition compared to compared to high-water availability when aggregated over effect sizes of all treatments.

## Discussion

The impact of increasing drought in a changing climate [60] and ever-present competition have large ramifications for natural plant communities and agricultural systems. Specifically, competition and water stress impacts community membership [3,61] and crop yield [10,62] and has global importance for plant conservation and food security. We demonstrate that water availability significantly modulates competitive outcomes where high-water availability intensified shoot-only competition while weakening root-only competition and the opposite patterns for low water availability. These results are important as short-term effects of competition were a top predictor of species’ abundance in the field [63]. This meta-analysis combines empirical evidence to reveal competitive patterns and influence future work to advance our knowledge.

### Shoot competition responses to water availability

We show that shoot-only competition was more intense under high-water availability than in low-water availability treatments. Higher aboveground biomass in high water availability treatments may have resulted from plentiful soil resources available for biomass production[27,29–31]. Furthermore, higher aboveground allocation could be in response to light competition for shade avoidance responses that can intensify competition through imposing shade [64,65]. From a community perspective, research suggests that light competition is important in ecosystems with high aboveground productivity [66] and thus aboveground competition can pattern community diversity and dynamics [67] and point to the importance of aboveground competition in different contexts.

To the contrary, the weakest competitive treatment was shoot-only competition in low-water availability. Water stress is known to limit plant growth leading to a reduction in leaf area which limits shading and light competition that an individual can impose on its neighbor [68]. These results seem to agree with the stress gradient hypothesis which notes that weaker competitive interactions may dominate at high-stress levels compared to low-stress [44,69]. Weaker competitive interactions could be a result of plants allocating less mass aboveground or slowing metabolic activity aboveground for survival and defense under stressful conditions [32]. This is interesting given that competition in dry environments is high, though thought to be concentrated belowground [70], however, we clearly demonstrate that when shoot competition is considered alone water availability is a key factor modulating its intensity and this needs exploration in different biomes.

### Root responses to water availability

Root-only competition was weaker at high-water availability. This suggests that higher water supply weakens belowground competition and shows different patterns to shoot-only competition. These results are in line with Lamb et al. [34], but counter Bartelheimer et al. [49] who showed competitive suppression in root-only treatments under high water supply. On the other hand, root-only competition was the most intense competition treatment under low-water availability. Intense root competition may be driven by roots responding to water stress by increasing root allocation and intensity of soil exploration resulting in increased nutrients and water uptake [24,25,71,72]. High root biomass and root length production are known to induce strong level of competition between plants [73] and these morphological changes in response to water stress likely increase the competitive environment as well [10,74]. Research suggests that root competition is more intense in dry environments where productivity is concentrated belowground [70,75]. Root-only competition was more intense than shoot-only competition under low-water availability. These results along with findings of Kiaer et al. [20] on nutrients indicate that when soil resources are limited, root competition is more intense than shoot competition. Despite this strong evidence of a positive effect of water shortage on the root competition we may expect conflicting responses when species evolve in differing environments, though more studies are needed to better assess this hypothesis.

### Whole plant outcomes and implications

The contrasting results between shoot-only, root-only, and full competition suggest that the contributions of root and shoot competition are not additive. Rajaniemi et al. [38] showed that root-only competition experimental assemblages resulted in lower species diversity compared to shoot-only competition assemblages. Also, Lamb et al. [76] showed shoot competition negatively impacted community evenness but was through indirect increases in competitive root responses. While aboveground competition has documented impacts on community structure [77], root competition also has strong and apparent consequences for plant communities.

Given the climate change outcomes of increased drought leading to increased root allocation [25] this may have important competition-mediated community outcomes. We may see increases in root competition for water in communities (sensu [33], [25]) that lead to plant diversity loss from drought [78]. But more research is needed to assess these outcomes and in different biomes. Because we see contrasting outcomes in root-only and shoot-only competition, researchers should increase the assessment of belowground ecology to draw more accurate conclusions about competition particularly if environmental constraints would lead to a shift in biomass allocation [79].

### Review of suitable studies not included in analyses

Two studies with suitable experimental designed lacked information for use in meta-analysis, but their results provide valuable information to the study outcomes. Semere & Froud-Williams [68] competed two pea cultivars with maize and found that pea cultivar identity and low-water availability impacted root-only competition. Both pea cultivar’s growth were not significantly affected by shoot-only competition, while root-only competition and low water availability reduced mass by 43%. These results indicate that root-only competition impacted growth while shoot-competition had smaller effects. Another study [80] competed adult *Lolium perenne* with *Phleum pratense* and *Trifolium pretense* seedlings. P. pretense was more affected by shoot-only competition than root-competition and shoot-competition was not modulated by water stress. Both species had higher root allocation at low water availability and under root-only competition. Root competition increased root allocation but not leaf area. Additionally, shoot-only competition was weakest at low-water availability while root-only competition was higher in low-water stress. The results of both studies highlight the variability in plant response to low-water availability and are in line with the findings of this meta-analysis that root-only competition is more intense than shoot-only competition.

### Study limitations

These results show important interactions between plant competition and water availability. The fixed effects used in these models significantly explained variation in effect sizes but including other effects such as target species life history, non-target life-history, and experimental setting may reduce residual heterogeneity. Given the small number of studies, these factors could not be reliably tested without replication. Other sources of variation were in the differences in materials used to partition plants (e.g. mesh vs. solid aboveground dividers) and implementation of water stress where amounts that were considered “high” and “low” differed by study. Additionally, the adaptations of target species could have influenced competitive outcomes and responses to water stress. For example, Bartelheimer et al. [49] used *Senecio aquaticus* – a wetland adapted species – which performed poorer than the terrestrial species in low water availability. Finally, there were known suitable studies that we excluded due to missing information and missing suitable studies introduce publication bias [81]. The results of relevant treatments in suitable studies were likely not reported due to lack of significance, introducing selective reporting bias [81]. Authors should publish robust study results and parameters (e.g. sample size, responses, measures of variability) for future synthesis and knowledge advancement.

## Conclusions

The intensity of root-only and shoot-only competition showed opposing trends under differing water availability. Our results show that roots have major implication in competitive outcomes for plants when soil resource are limited. This suggests that root-dominated interactions should make coexistence more difficult and lead to more growth suppression in case of water shortage. Importantly, if we only record aboveground responses to water stress or competition we may conclude weak competition when belowground responses may reveal contrasting evidence. Future research should tie in the role that root and shoot competition has on species coexistence in plant communities.

## Acknowledgements

I extend gratitude to Dr. Larry Hedges, PhD for invaluable feedback on early manuscript versions and Dr. Holly Jones for feedback this manuscript. I thank Dr. Alexandra Weigelt, PhD for providing annotated and translated data from her study for inclusion here. I thank Dr. Andrea Kramer, PhD and the Kramer-Havens lab group for suggestions on an early version of this document. Finally, I thank Drs. Amy Iler, PhD and Stuart Wagenius, PhD for input on analyses.

---

**Supplemental Figure 1. PRISMA flow diagram**. Peer-reviewed journal articles were searched in Web of Science. Key-terms are: [(shoot* AND root*) OR (above AND below)] AND [(competit* OR interact*)], topic: “water stress.”

**Supplemental Table 1. Study Data.** Study dataset used to calculate effect sizes and sampling variances.

**Supplemental Table 2. Checklist.** PRISMA checklist.

